# Disease-associated protein states accessible to 8-oxoguanine transcriptional miscoding without genomic mutation

**DOI:** 10.64898/2026.10.01.756037

**Authors:** Santiago Moreno-Caceres, Alexis M. Hayes, Amber Delmas Eliason, David P. Marciano

## Abstract

8-Oxoguanine (8-oxoG) is a common oxidative DNA lesion that can alter coding output during transcription without changing the underlying DNA sequence. When present on the template strand, 8-oxoG can direct RNA polymerase II to insert adenine, generating a transient coding-sense C>A error restricted by the genetic code to 18 directional amino-acid substitutions. Mapping these substitutions onto ClinVar identified 3,910 disease-associated protein states across 1,333 genes, representing 5.7% of evaluable pathogenic or likely pathogenic missense states. In 211 cases, transcriptional miscoding and the corresponding genomic disease variant affect different nucleotides within the same codon yet converge on the same amino acid substitution, allowing the disease-associated protein state to arise while the implicated genomic nucleotide remains wild type. Accessible states were enriched among genes with post-mitotic expression and neurological disease annotation, particularly in neuronal signaling pathways. Published oxidized-guanine maps supported 2,146 accessible genomic coordinates, while human proteogenomic data contained 78 exact codon-compatible protein states not explained by sample-specific transcript sequence. Together, these analyses define a chemically constrained set of disease-associated protein states accessible through 8-oxoG transcriptional miscoding without permanent genomic mutation.

## Introduction

Protein states associated with genetic disease are usually defined by genomic variants, yet the same altered amino-acid sequence can sometimes arise without a permanent mutation. DNA lesions can transiently change the sequence copied into RNA during transcription while leaving the underlying genomic sequence unchanged. Template-strand 8-oxoguanine provides a chemically defined example, because RNA polymerase II can insert adenine opposite the lesion and produce a coding-sense C>A error in the transcript.

The biological consequence of 8-oxoguanine depends on which polymerase encounters the lesion. During DNA replication, inaccurate base incorporation due to guanine oxidation can convert the lesion from a permanent G:C to T:A substitution at the oxidized base pair [1], contributing to the oxidative mutational signatures observed in human cells and cancers [2, 3]. During transcription, the same lesion instead alters only the nascent RNA, and repair determines how long the template remains available for repeated miscoding [4, 5]. Structural studies of the RNA polymerase II elongation complex explain this dual coding potential, because 8- oxoguanine in the *anti* conformation pairs with cytosine, whereas rotation to the *syn* conformation permits a Hoogsteen-like interaction with adenine [6]. Polymerase bypass of the lesion has been characterized directly [7, 8], mammalian reporter assays have shown that defined 8-oxoguanine generates transcriptional errors at measurable frequencies [9], and error-corrected sequencing has independently quantified rare transcription errors in human cells [10]. A defined template lesion at HRAS codon 61 produced a repair-dependent signaling response [11], and transcriptional mutagenesis of alpha-synuclein has been reported in Parkinson’s disease [12].

Throughout this work, we define a disease-associated protein state by an amino-acid substitution encoded by a ClinVar pathogenic or likely pathogenic missense variant, alongside a higher-confidence subset defined in Methods. The miscoding chemistry of 8-oxoguanine is established, but the extent to which its genetically constrained amino-acid outcomes intersect the protein states that human genetics implicates in disease remains undefined. Because the coding consequences of a template-strand 8-oxoguanine are fixed by the genetic code, we can define the full missense repertoire of this route without assuming how frequently any individual lesion occurs. We mapped that repertoire onto pathogenic and likely pathogenic human variation, cell-expression context, disease annotation, published oxidized-guanine maps, and directly observed protein substitutions.

## Results

### The genetic code confines 8-oxoguanine miscoding to 18 directional amino-acid substitutions

When RNA polymerase II inserts adenine opposite a template-strand 8-oxoguanine, the resulting transcript carries a coding-sense C>A error whose protein consequence is determined by the genetic code (Figure 1a). Applying a coding-sense C>A change to every cytosine in every sense codon yields 48 codon-position combinations: 33 missense, 11 synonymous, and 4 stop codons. The 33 missense outcomes collapse to 18 distinct directional amino-acid substitutions because several codons for the same residue yield the same product (Figure 1b).

**Figure 1.**
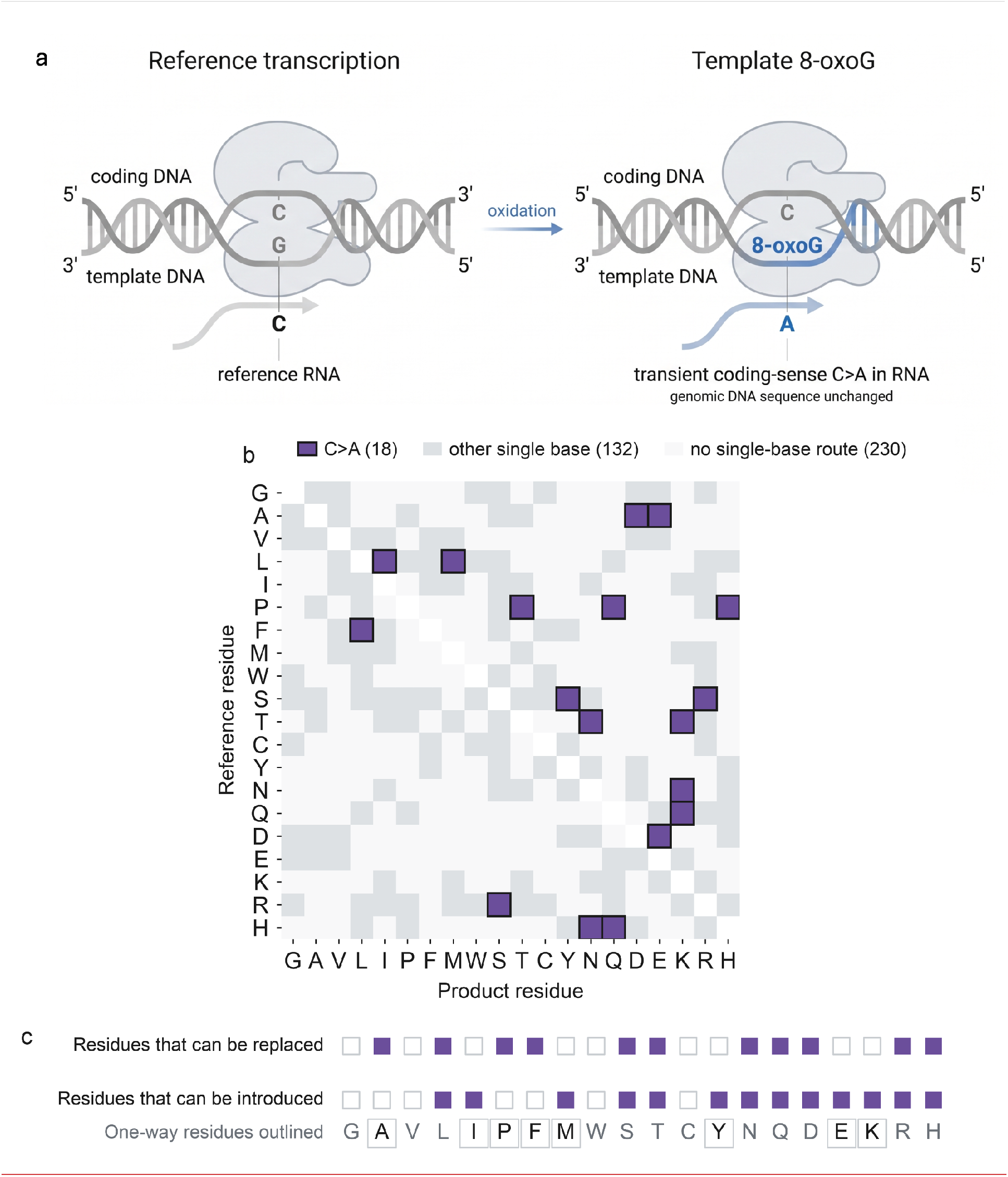
The genetic code confines 8-oxoguanine transcriptional miscoding to 18 amino acid substitutions. (a) Base pairing at the polymerase active site, where template guanine pairs with cytosine through Watson-Crick geometry, while 8-oxoguanine can rotate to the syn conformation and pair with adenine through Hoogsteen-like geometry, giving a coding-sense C>A change in the transcript, with hydrogen bonds dashed and geometry following published RNA polymerase II elongation-complex structures [6]. (b) All ordered amino-acid pairs, with purple marking the 18 pairs reachable by a coding-sense C>A change, gray the 132 reachable by another single-base change and pale gray the 230 requiring more than one. (c) Residues this route can replace and can introduce, where outlined codes mark the eight residues accessible in one direction only. Figure 1a created with BioRender https://BioRender.com/1j03jso.

Because the outcome is constrained by both lesion chemistry and the genetic code, the route permits some protein states while making others inaccessible regardless of lesion frequency. This route can replace 11 residues and introduce 13, with eight of the 20 amino acids accessible in only one direction (Figure 1c and Supplementary Table S1). Glycine makes the constraint especially clear, because every coding-sense C>A change in a glycine codon is synonymous, so glycine substitutions are inaccessible regardless of how frequently the lesion occurs. Tyrosine cannot be replaced because the corresponding change produces a stop codon, although serine can be introduced as tyrosine.

### Accessible substitutions intersect thousands of disease-associated protein states

Despite this narrow chemical route, the 18 accessible substitutions are compatible with thousands of protein states already associated with human disease. Intersecting the accessible substitution classes with ClinVar pathogenic and likely pathogenic missense records mapped to MANE Select transcripts [13, 14] gives 3,910 disease-associated protein states in 1,333 genes (Supplementary Table S2). These states represent 5.7% of all pathogenic or likely pathogenic ClinVar missense states that reconcile to the analyzed MANE Select transcripts. Restricting the analysis to variants with at least two ClinVar review stars and excluding splice-proximal positions retained 977 states across 403 genes. The states are distributed across all 18 substitution classes rather than concentrated in one (Figure 2a), and most are reachable by a genomic variant at the same nucleotide, with the remainder split between the two convergence classes (Figure 2c). Thus, a chemically restricted transcriptional error can access thousands of protein states already implicated by human disease genetics.

**Figure 2.**
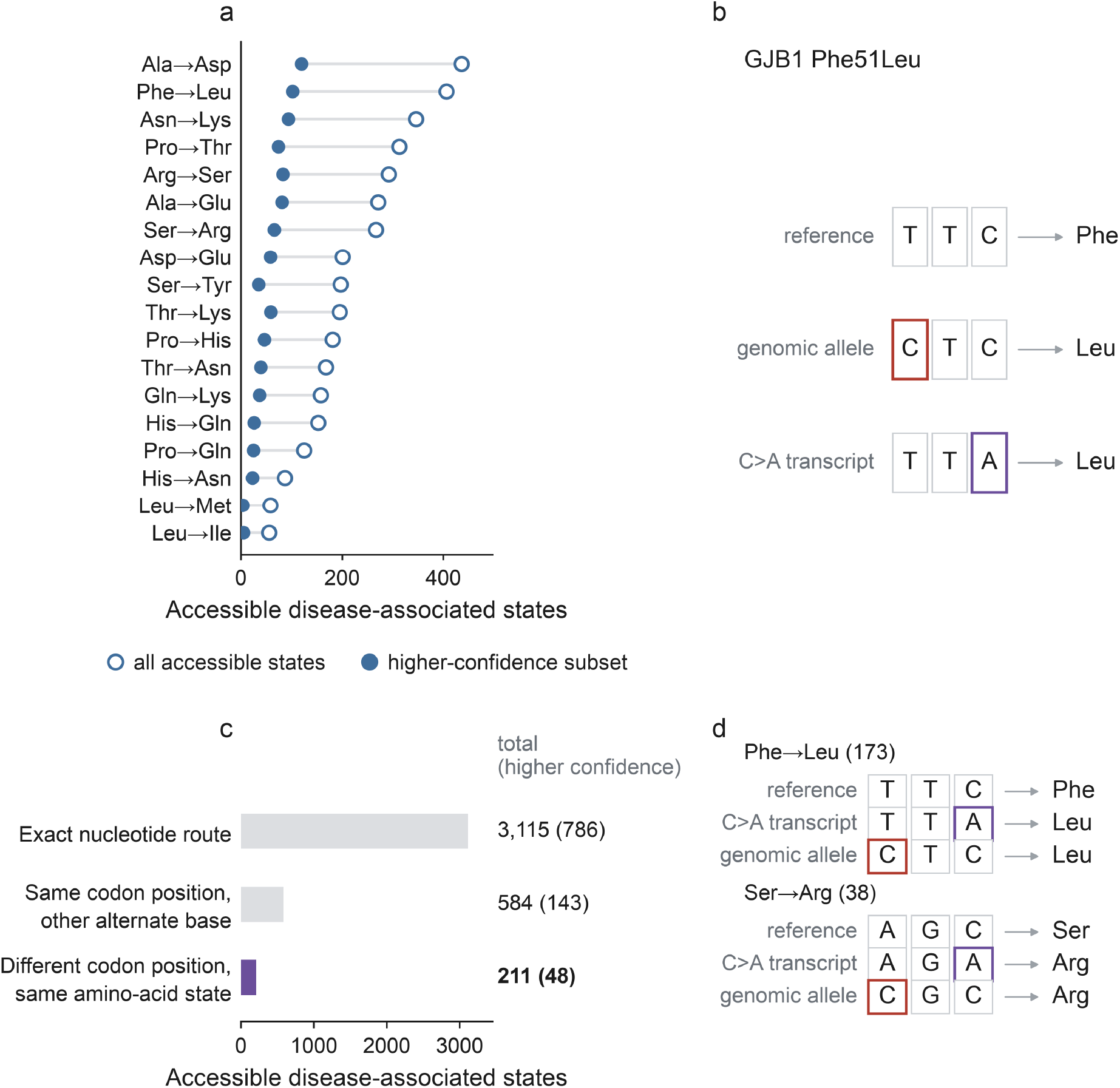
Accessible substitutions intersect 3,910 disease-associated protein states, including 211 that can arise while the disease-associated nucleotide remains wild type. (a) States per substitution class, with open blue circles for all accessible states and filled blue circles for the higher-confidence subset. (b) GJB1 phenylalanine 51 as a worked example, where red marks the nucleotide altered by the pathogenic genomic allele and purple the nucleotide altered by the C>A transcriptional route, both producing leucine. (c) States by the relationship between the genomic and transcriptional routes within the codon, with purple marking the different-position class and values given as totals with the higher-confidence subset in parentheses. (d) The two codon configurations that account for all different-position states, with red and purple as in panel b.

### The same protein state can arise while the disease-associated nucleotide remains wild type

In 211 states, the coding-sense C>A error and the pathogenic ClinVar variant alter different nucleotides in the same codon but produce the same amino-acid substitution, so the disease-associated protein state can arise while the nucleotide implicated by human genetics remains wild type. These 211 states span 172 genes, and 48 belong to the higher-confidence subset. In each case, the transcriptional change occurs at the third codon position while the genomic variant alters the first, and two codon configurations account for all of them: phenylalanine to leucine from TTC and serine to arginine from AGC (Figure 2d). In GJB1, which causes Charcot-Marie-Tooth disease type X1, the pathogenic variant at phenylalanine 51 changes the first position of the TTC codon, while a coding-sense C>A change at the third position produces the same leucine substitution. In that case, transcriptional miscoding could produce the disease-associated protein state while sequencing of the position implicated by human genetics returns the reference base (Figure 2b). This class separates the amino-acid state from the nucleotide through which it was identified.

### Accessibility is associated with combined post-mitotic expression and neurological disease annotation

Chemically accessible disease-associated states were not distributed uniformly across gene contexts. We interpreted these differences as associations with gene annotation rather than as evidence that lesions form, persist, or miscode at different rates across cell types. Across all eight single-cell expression classes of the Human Protein Atlas [15], neurons are the only class in which the accessible fraction exceeds its gene-label permutation expectation, at 7.67% against 6.03% (q = 0.004, Figure 3a). The other post-mitotic classes fall inside their permutation intervals, and oligodendrocytes fall below, so this is not a general property of post-mitotic tissue. Across all 10 top-level MONDO disease categories [16], nervous system disease had the highest accessible fraction, although the gene-level screen did not survive correction (q = 0.21, Supplementary Table S3a).

**Figure 3.**
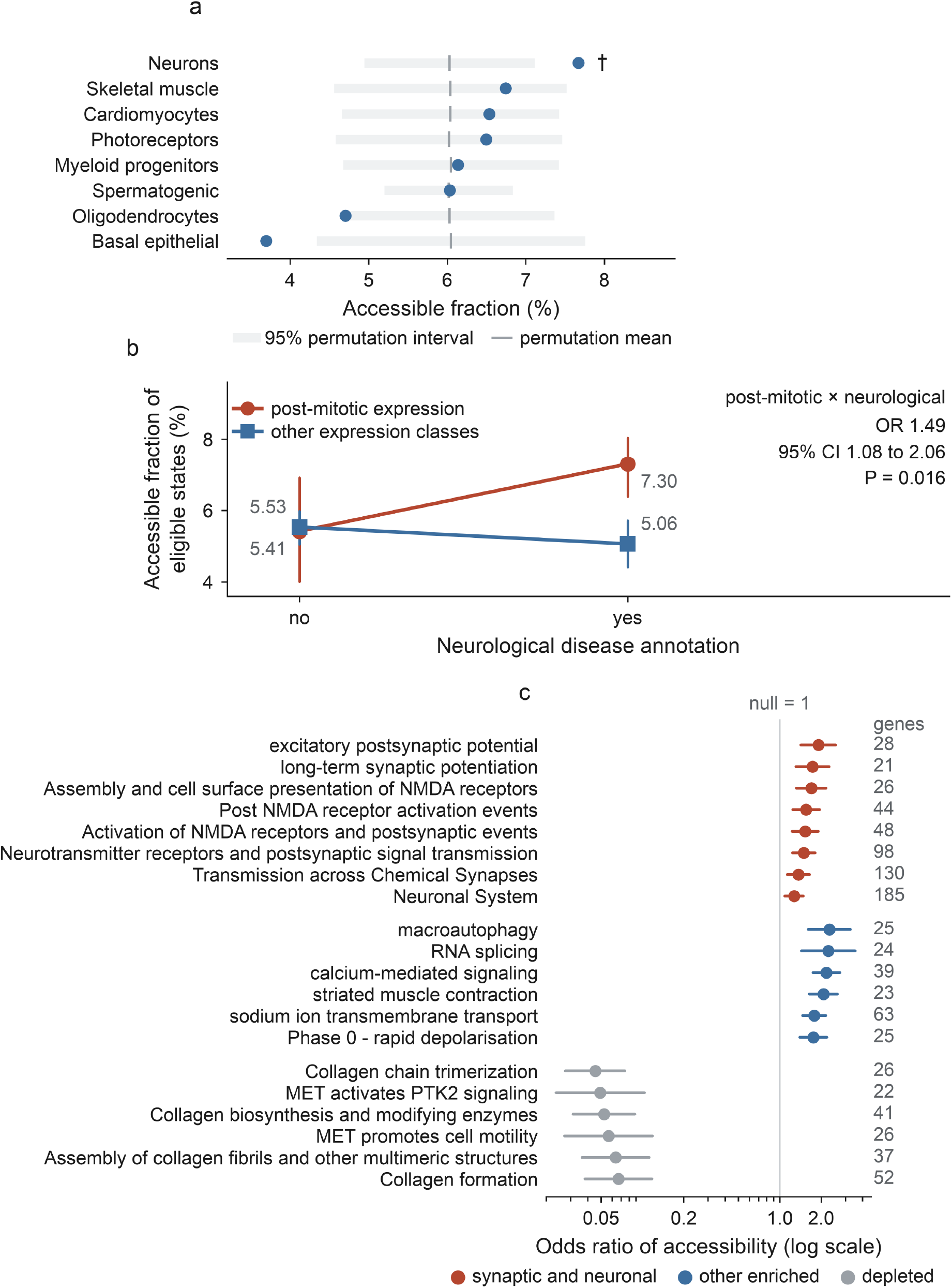
Chemically accessible disease-associated states vary with combined post-mitotic expression and neurological disease annotation. (a) Accessible fractions across eight Human Protein Atlas single-cell expression classes, with blue points showing observed values, gray bars the 95% gene-label permutation intervals and gray ticks the permutation means, where neurons were the only class surviving correction (†q = 0.004). (b) Accessible fractions jointly stratified by post-mitotic expression and neurological disease annotation, with whole-gene bootstrap intervals and the interaction estimate of the joint gene-level model shown on the panel. (c) Pathway associations with 95% confidence intervals on a logarithmic odds-ratio axis, colored by family, where every displayed pathway passed a q-value below 0.05 and was selected by the prespecified rule described in Methods.

The combination of post-mitotic expression and neurological disease annotation carried more accessible states than either annotation alone would predict. In a gene-level model of accessible states among eligible states per gene, adjusted for intrinsic C>A coding opportunity, coding length, and ClinVar record count, post-mitotic expression alone showed no independent association (odds ratio 0.98; 95% CI 0.74 to 1.29). The interaction between post-mitotic expression and neurological annotation corresponded to an odds ratio of 1.49 (95% CI 1.08 to 2.06, P = 0.016, Figure 3b). The signal was associated with the combination of post-mitotic expression and neurological disease annotation rather than with post-mitotic expression alone, and it holds across alternative specifications, including exclusion of neuron-enriched genes and restriction to the higher-confidence subset (Supplementary Figure S1). We constructed the expression classification after the accessible-state catalog was complete, so we report this analysis as exploratory rather than as a test of a prespecified hypothesis. The association identifies a gene context in which chemically accessible states are more common without implying that neurons carry more lesions, repair them less efficiently, or miscode more frequently.

Pathway analysis pointed to the same biological context. Among 945 estimable gene sets, 102 passed a false-discovery threshold of 0.05, of which 64 were enriched and 38 depleted (Figure 3c and Supplementary Table S4). Of the 30 sets annotated to synaptic or neuronal signaling, the eight that passed correction were all enriched; these included excitatory postsynaptic potential, long-term synaptic potentiation, and the NMDA receptor and neurotransmitter receptor pathways. Enriched sets outside that group include calcium-mediated signaling and sodium ion transmembrane transport. Four of the six most strongly depleted sets involve collagen or extracellular-matrix biology, a pattern consistent with the inaccessibility of glycine substitutions through the C>A route.

### Published oxidized-guanine maps detect many accessible disease-associated coordinates

We asked separately whether accessible disease-associated coordinates have been detected in oxidized-guanine maps and whether they are detected more often than comparable coding positions. Published maps detected oxidized-guanine signal at 2,146 of the 3,910 accessible disease-associated coordinates, including 559 of the 977 higher-confidence positions, providing direct mapping support for many chemically accessible coordinates (Supplementary Figure S2 and Supplementary Table S6). Detection was similarly common among route-matched missense-capable coding cytosines in the same transcripts, where 346,415 of 638,965 background coordinates were detected. Accessible disease-associated positions were not preferentially detected in these oxidized-guanine maps, at an odds ratio of 1.03 (95% CI 0.96 to 1.09, P = 0.40) [17, 18]. Restricting the gene-context model to map-supported coordinates preserved the post-mitotic-by-neurological interaction and strengthened it, yielding an odds ratio of 1.77 (95% CI 1.22 to 2.57, P = 0.0027) versus 1.49 in the primary model. The whole-gene bootstrap interval was 1.20 to 2.60, and the estimate in the higher-confidence subset was 2.87 (95% CI 1.42 to 5.80) (Supplementary Figure S1). Map detection within the accessible states was not associated with post-mitotic expression, neurological annotation, or their interaction after adjustment for coding and sequence covariates. The persistence of the gene-context association among map-supported coordinates argues against preferential detection in that gene context as the cause (Supplementary Table S6). A coordinate without map support is not established as lesion-free because detection depends on assay sensitivity, the cells, and the conditions each resource was sampled under.

### C>A-accessible protein states occur in human tissues among substitutions unexplained by sample-specific transcript sequence

To determine whether chemically accessible states are observed in human proteins, we queried a published proteogenomic survey of more than 1,000 human samples. In that survey, we searched each sample against a protein database built from its own RNA sequencing, including the variants called in that sample, so substitutions explained by sample-specific transcript sequence were excluded by construction [19]. Of 1,955 confidently localized substitution sites, 1,658 reconciled to MANE Select transcripts, 93 fall in a substitution class this chemistry can produce, and 78 are reachable at the exact codon position (Figure 4a). The 78 states occur across 44 genes, with 25 in three hemoglobin genes, consistent with the greater detectability of highly abundant proteins (Figure 4b and Figure 4c). The source study separately analyzed the matched whole-exome sequencing available for the cancer cohorts and detected a missense variant capable of encoding the observed substitution in only two cases. It attributed the remaining events to non-genetic decoding mechanisms, including alternative translational decoding and RNA modification [19]. The 78 exact codon-compatible states demonstrate that amino-acid outcomes available to the C>A route occur in human tissues, although the proteogenomic data do not identify the molecular process responsible for any individual event.

**Figure 4.**
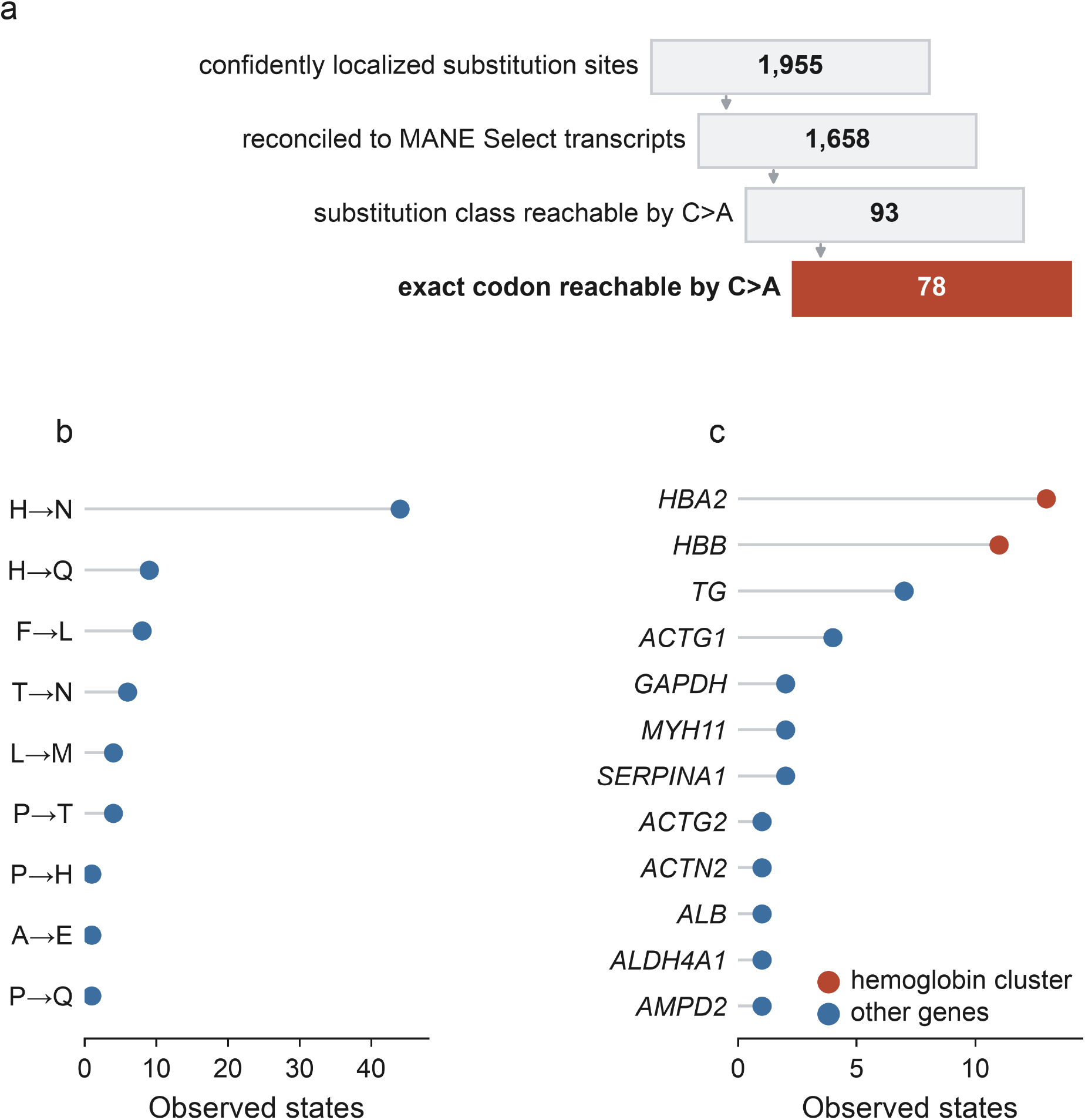
C>A-accessible protein states are observed in human tissues among substitutions unexplained by sample-specific transcript sequence. (a) Stepped filtering cascade from confidently localized substitution sites to the states reachable at the exact codon position, with the count printed inside each step, uniform step widths that do not encode those counts and the accent color marking only the final group. (b) The 78 accessible states by substitution class. (c) The 12 genes with the most observed accessible states, ranked by count and then alphabetically, with hemoglobin genes highlighted.

## Discussion

A disease-associated protein state does not require the pathogenic nucleotide that first identified it. One-directional transcription error, confined by the genetic code to 18 amino-acid substitutions, reaches 3,910 such states across 1,333 genes. The 211 different-position states make the separation clearest, because the genomic disease allele and the transient transcriptional error alter different nucleotides of one codon while converging on the same residue, so the protein state can be present while the position human genetics implicates as reference. Human genetics has established disease associations for variants encoding these protein states, providing a biologically grounded set of amino-acid outcomes in which to examine the potential consequences of transient transcriptional miscoding. A transiently produced state would not reproduce the dosage, tissue distribution, or developmental timing of an inherited variant, but it would reproduce the amino-acid sequence through which that variant acts.

Long-lived post-mitotic cells provide a distinctive biological setting for transcriptional miscoding. Replication is minimal or absent while transcription persists, so a lesion that escapes repair can affect multiple transcripts without becoming fixed in the genome, with repair kinetics rather than replication determining how long that opportunity remains. Transcriptional mutagenesis of alpha-synuclein has been reported in Parkinson’s disease [12], which shows the phenomenon is detectable in neurological tissue. Our gene-context result identifies where chemically accessible states are concentrated rather than where lesions occur or where repair fails.

The strongest direct precedent remains a defined template-strand 8-oxoguanine at HRAS codon 61, which produced increased ERK phosphorylation in repair-deficient mouse fibroblasts and not in repair-competent cells [11]. That experiment established that a single defined lesion can change cellular signaling through transcription, whereas the present analysis defines the protein-state space this chemistry could access across human disease genetics. Because the sequence change first appears in RNA, transcriptional miscoding can also affect RNA processing before translation, consistent with its reported effects on splicing fidelity [20]. The consequences of a miscoding event may extend beyond the amino-acid substitution itself.

Several limitations define what this landscape can establish. Chemical accessibility describes which amino-acid outcomes the genetic code permits through this route, but it does not establish how often any occur. Whether a particular state is produced depends on lesion formation on the template strand, the lesion’s position within the transcribed sequence, transcription of that allele, repair kinetics, RNA turnover, translation, and the stability of the resulting protein, and none of these quantities is measured here. The published oxidized-guanine maps used in this work report lesion positions in the cells and conditions those studies sampled, which does not establish lesion occupancy in the tissue or cell type relevant to any particular disease. The proteogenomic observations demonstrate that amino-acid outcomes compatible with this chemistry occur in human tissue, and they do not attribute those outcomes to 8-oxoguanine. A transient low abundance proteoform should also not be assumed to reproduce the dosage, timing, or tissue distribution of an inherited pathogenic allele. This map defines what the genetic code and lesion chemistry permit, but it does not establish that any individual state is produced at a consequential level or contributes to disease.

By defining the complete directional landscape imposed by 8-oxoguanine miscoding, this study moves transcriptional mutagenesis from isolated examples toward a genome-scale framework for identifying the disease-associated protein states available to oxidative DNA damage. Which of those states are produced will depend on lesion position, transcription, repair, and protein stability, making oxidative stress a potential modifier of the proteoform repertoire produced by a cell without requiring permanent mutation. The critical experimental step is to link a defined template 8-oxoguanine to its RNA error and predicted protein state in the same biological system, while confirming that the genomic sequence remains unchanged.

## Methods

### Transcripts and coding sequences

We obtained coding sequences and transcript structures from MANE Select v1.5 [13], which contains 19,367 protein-coding records. We retained the 19,275 transcripts whose coding sequence begins at an annotated start codon, ends at the first in-frame stop, and has a length matching the summed genomic coding blocks to within one codon. The 92 excluded records are itemized with their exclusion reason in the transcript quality-control record distributed with the reproducibility archive, and none of them contribute to the main-text results. Protein coordinates refer to the retained transcript, and every record used in an analysis had to translate to its annotated wild-type residue.

### Accessible substitution enumeration

For every sense codon and every cytosine within it, we replaced the cytosine with adenine and translated the product, classifying outcomes as missense, synonymous, or stop-gained, and collapsed amino-acid pairs across codons to define substitution types. Replaceability is the fraction of a residue’s synonymous codons admitting a missense change, and introduction potential is whether any codon of another residue yields it as a product. The lesion sits on the template strand, so a template guanine corresponds to a coding-sense cytosine, and the transcript change is C>A.

### Clinical variants

We retrieved variant records from the ClinVar GRCh38 variant_summary.txt.gz release on August 27, 2026 [14]. We retained records with an aggregate ClinVar clinical significance of Pathogenic, Likely pathogenic, Pathogenic/Likely pathogenic, or another pathogenic classification with no conflicting, uncertain, or benign interpretation. A record also required a codon in the MANE transcript translating to the annotated reference residue. We mapped each qualifying single-nucleotide variant to its MANE codon and codon position for route classification, then collapsed records to one row per gene, protein position, and amino-acid substitution for the state-level analyses. The higher-confidence subset required at least two ClinVar review stars, corresponding to multiple submitters without conflicts, expert-panel review or a practice guideline, and excluded coding positions within three nucleotides of an annotated exon boundary.

### Alternative nucleotide routes to disease-associated amino-acid states

For each accessible missense state, we recorded the wild-type MANE codon, the position of the accessible cytosine, the resulting codon, and its amino-acid product. We mapped pathogenic and likely pathogenic single-nucleotide ClinVar variants producing the same amino-acid substitution at the same protein coordinate back to that codon. We defined three mutually exclusive classes based on how the documented genomic allele relates to the C>A transcriptional event within the codon. Exact nucleotide concordance alters the same nucleotide to the same base; same-position alternate-base convergence alters the same nucleotide to a different base; and different-position convergence alters a different nucleotide of the codon, the last two producing the same amino-acid substitution by a different nucleotide change. The three classes contain 3,115, 584 and 211 records and sum to the 3,910 classified states. We excluded multi-nucleotide variants and applied the codon reconciliation requirement described under Clinical variants without modification.

### Experimentally mapped oxidation positions

We retrieved two published resources mapping oxidatively modified guanine at fixed releases before analysis, a base-resolution map from human cells [17] (GSE181312) and click-code-seq position lists [18] (GSE295218). We applied the same mapping procedure to both, and each resource is identified by accession, release, and retrieval date in the source manifest distributed with the reproducibility archive. Signal thresholds are not detection guarantees, so we use both maps as positive coordinate annotations, and we do not classify an undetected position as lesion-free. Click-code-seq is strand-resolved, so support required detection on the transcription template strand. The base-resolution map is not strand-split, so we retained detections as coordinate-level support at the genomic base pair without inferring which strand carried the lesion. We converted coordinates to the MANE transcript model on GRCh38, applying liftOver when a source deposit used GRCh37 and support was required at a coding position. Both assays primarily detect 8-oxoG but can also detect FapyG at lower abundance. We scored the lesion-detection rate against one route-matched background comprising all 638,965 missense-capable coding cytosines in the 1,333 retained transcripts that carry at least one accessible disease-associated state, excluding the accessible positions themselves, and tested it with a Fisher exact test.

### Human proteogenomic state mapping

We obtained directly observed noncanonical protein states from the substituted-peptide tables distributed with the published proteogenomic survey of more than 1,000 human samples across 26 healthy tissues and six cancers [19]. The source study’s search design excludes substituted peptides explained by the sample’s own transcript sequence, and the Supplementary Methods describe that design and the basis on which the source attributed the remaining events to non-genetic decoding. We retained states only when the source assigns the substitution to a single residue, yielding 1,955 confidently localized sites out of the 8,746 reported. Each site was reconciled to MANE Select v1.5 by requiring exact agreement of gene symbol, protein position and reference residue, leaving 1,658 evaluable states. We called a state exactly C>A-accessible when replacing one specific cytosine in its actual MANE codon with adenine produces the observed residue. This codon-level test is stricter than membership of an accessible substitution class and removed 15 of the 93 class-compatible sites, leaving 78. For each accessible state, we converted the coding cytosine whose change produces the observed residue to a genomic coordinate and resolved the corresponding template-strand guanine, which succeeded for 77 of the 78 states; we queried both map resources there. We called support endogenous when the coordinate was detected without exogenous oxidant and induced-only when detection occurred solely under induced conditions. Sensitivity analyses removed the hemoglobin gene cluster and the histidine-to-asparagine class, separately and together, because both dominate the observed set.

### Pathway analysis

We obtained gene sets from Reactome [21], using the release recorded in the source manifest, and from Gene Ontology biological process annotations [22] in the GOA human release of 26 July 2026. We read term names from the GO-BASIC ontology, confirmed that every identifier lies in the biological process namespace, and mapped NCBI gene identifiers to symbols using the NCBI gene information file. We used the gene-context model universe for testing and retained sets with 20-500 universe genes. For each set, we modeled the per-gene count of accessible states out of eligible states using binomial logistic regression, with set membership as the predictor and standardized C>A coding opportunity, log ClinVar record count, and log coding length as covariates. Because amino-acid composition directly constrains accessibility to a coding-sense C>A change, the models additionally included the standardized fraction of protein residues replaceable by this route, computed from the MANE Select protein sequences. Wald inference used heteroskedasticity-robust HC1 standard errors, and dispersion in the covariate-only model was 1.07. Sets in which no gene carried an accessible state are not estimable and are excluded, and false-discovery correction used the Benjamini-Hochberg procedure across the estimated sets. Figure 3c displays a prespecified subset of the sets passing correction, comprising every set annotated to synaptic or neuronal signaling, the six enriched sets outside that group with the largest odds ratios, and the six depleted sets with the smallest odds ratios. We annotated a set as synaptic or neuronal signaling if its name contains synapse, neurotransmitter, NMDA, neuron, axon, dendrite, glutamatergic, or GABAergic and does not also refer to DNA pairing, recombination, or meiosis; this excludes two DNA-repair sets whose names contain the word synapsis in an unrelated sense. Supplementary Table S4 reports every estimated set, the displayed subset, and the reason each displayed set was selected.

### Software

Analyses used Python 3.13, pandas 2.3.3, numpy 2.3.4, scipy 1.16.2, statsmodels 0.14.6, matplotlib 3.10.7, openpyxl 3.1.5, and pillow 12.0.0.

### Data and code availability

All primary resources analyzed in this study are publicly available and are identified by accession, release and retrieval date in the source manifest. Variant records are from ClinVar, reference transcripts and proteins from MANE Select v1.5, expression classes from the Human Protein Atlas single-cell type consensus data, disease categories from MONDO through ClinVar phenotype terms, gene sets from Reactome and the Gene Ontology annotation, oxidized-guanine coordinates from GSE181312 and GSE295218, and the directly observed protein states from the supplementary source data of the cited proteogenomic survey. The derived data supporting the figures and reported values are provided in Supplementary_Tables.xlsx and as machine-readable files in the reproducibility repository. The repository contains the frozen analysis inputs, the analysis and figure code and the pinned software environment required to rebuild the Supplementary Tables and figures and to recompute the reported numerical results that can be checked from those files. Six quantities, which are published counts, extraction intermediates or the route-matched background comparison, are not recomputed from the distributed files. The eight expression classes in Figure 3a and the ten disease categories regenerate from frozen per-class records. Primary source files are not redistributed. Code and derived data are available at https://github.com/Marciano-Lab/8oxoG-disease-transcriptional-miscoding.

## Declarations

### Ethics approval and consent to participate

Not applicable. This study analyzed only publicly available, previously published datasets and curated database records.

### Consent for publication

Not applicable.

### Competing interests

The authors declare no competing interests.

### Funding

This work was supported by the National Institutes of Health, National Heart, Lung, and Blood Institute (R00HL145097) and the American Heart Association (23IPA1053651 and 26BIPA1622618) to D.P.M.

### Authors’ contributions

All authors performed the analyses. D.P.M. wrote the manuscript, and all authors edited and reviewed the final version.

## Notes

https://github.com/Marciano-Lab/8oxoG-disease-transcriptional-miscoding

### Competing Interest Statement

The authors have declared no competing interest.

