## Supplemental_Information for "Disease-associated protein states accessible to 8-oxoguanine transcriptional miscoding without genomic mutation"

*References are numbered as in the main manuscript.*

### Supplementary Methods

#### ClinVar route classification

We performed route classification on variant-level records before collapsing to amino-acid states, preserving the nucleotide identity needed to distinguish exact, same-position alternate-base and different-position routes. We excluded multi-nucleotide variants and records that failed exact reconciliation to the MANE Select coding sequence; in the final qualifying set, each of the 3,910 retained states was represented by exactly one qualifying single-nucleotide variant, so the three route classes partition the state set directly. No state was added manually. We independently audited the completeness of the frozen ClinVar extraction against later archived records and by codon-level testing of candidate accessible substitution classes, and we include the full audit in the reproducibility archive.

#### Oxidized-guanine map coordinate conversion

Two published resources contributed coordinates. Base-resolution coordinates were taken from the three wild-type HeLa samples and the four oxidant-exposed in vitro and in vivo samples deposited under GSE181312, which provide the endogenous and induced condition categories, respectively. We obtained strand-resolved Click-code-seq coordinates from the twelve cellular samples deposited under GSE295218, comprising HAP1 and U2OS cells exposed to vehicle or the DNA methyltransferase inhibitor GSK-3484862 in three biological replicates each; we did not use the four plasmid control samples from that series. The same publication deposited a separate series (GSE272366), which we did not use.

Both map resources were converted to one coordinate convention on GRCh38 before intersection. We converted deposits released on GRCh37 using liftOver with the UCSC chain file, and discarded positions that failed to lift or mapped to more than one location. Converted positions had to fall within the coding region of a retained MANE transcript. We used published signal thresholds and did not re-derive any here, so we record a coordinate as detected or not detected under the source definition. Per-sample retained-coordinate counts, the per-step filter audit and the strand provenance of each resource are provided in the reproducibility archive.

**Click-code-seq coordinate constraint**

Click-code-seq reports the position immediately 5' of the excised lesion rather than the lesion itself, so each deposited coordinate was shifted by one nucleotide in the direction defined by the reported strand before intersection, and coordinates whose shifted position left the coding sequence were discarded. Because the assay is strand-resolved, we counted support only when the shifted position fell on the transcription template strand, which carries the guanine whose oxidation produces a coding-sense C>A transcript.

**Lesion query for the directly observed proteomic states**

For each of the 78 exactly C>A-accessible observed states, we converted the coding cytosine whose substitution produced the observed residue to a genomic coordinate using the MANE transcript model, resolved the template-strand guanine from transcript orientation, and queried both contributing map resources at that coordinate. We classified a state as endogenous when its coordinate was detected in a dataset generated without exogenous oxidant; induced-only when detection occurred solely under bromate or another induced oxidative condition; not detected when the coordinate was evaluable but absent from all retained maps; and unresolved when the template-strand guanine coordinate could not be assigned. These four categories are mutually exclusive and are reported per state in Supplementary Table S5.

**Basis of the alternate-decoding attribution**

The proteogenomic source assembled each sample's own RNA sequencing, called variants from it, and translated those calls into a sample-specific protein database, so substituted peptides explained by sample-specific transcript sequence were excluded by construction. The source study separately analyzed the matched whole-exome sequencing available for the cancer cohorts and detected a missense variant capable of encoding the observed substitution in only two cases. On that basis, the source attributes the surviving events predominantly to alternate translational decoding rather than to nucleotide-level change. Accordingly, these states are used only as evidence that C>A-accessible amino-acid outcomes occur in human tissue and not as evidence for the mechanism that generated them.

**Reconciliation of the observed proteomic states to MANE Select**

Of the 1,955 confidently localized substituted-peptide sites, 1,658 reconciled to the MANE Select protein definition used throughout, requiring exact agreement of gene, protein position, and reference residue. Ninety-three of the reconciled sites belong to one of the 18 accessible substitution classes, and 78 of those satisfied the stricter requirement that the coding-sense C>A change fall at the exact codon position producing the observed residue. The reproducibility archive itemizes the categories of reconciliation failures.

**Composition of the directly observed accessible states**

The 78 exactly C>A-accessible states span 44 genes; HBA2 (13 states), HBB (11), TG (7), and ACTG1 (4) contribute nearly half, and histidine-to-asparagine accounts for 44 of the 78. Both features are expected in a peptide-level survey, where detection favors abundant proteins and particular substitutions, and both prevent a simple enrichment claim. Removing the hemoglobin cluster leaves 53 states in 41 genes; removing histidine-to-asparagine leaves 34 states in 25 genes; and removing both leaves 25 states in 23 genes, 12 of which carry mapped oxidized guanine. Two of the 78 states match a pathogenic or likely pathogenic ClinVar state exactly, HBB His93Asn and HBB His98Asn, both arising from a CAC codon through a first-position C>A event, and both coordinates carry experimentally mapped oxidized guanine. Neither reaches high ClinVar review status, so we describe them as examples of multi-layer convergence rather than genetic validation. Per-state values for all of these comparisons are given in Supplementary Table S5.

**Gene-context screens and model**

We assigned genes to a single-cell expression class using the Human Protein Atlas single-cell type consensus data [15] and to top-level disease categories through the MONDO ancestry of their ClinVar phenotype terms. We screened both annotations in full rather than restricting them to a preselected class or category. For the disease screen, we compared each category with all others, with the gene as the unit of analysis, and adjusted category P values across the ten categories using the Benjamini-Hochberg procedure; the expression-class permutation is described under Expression-class construction. Supplementary Table S3a reports both screens in full, including classes and categories with no signal.

Gene-level accessibility was then modeled as a binomial response, with accessible states as successes and eligible disease-associated states as trials, fitted by binomial logistic regression. Predictors were post-mitotic expression class, neurological disease annotation, their interaction, per-gene transcriptional opportunity, the natural logarithm of ClinVar record count, and the natural logarithm of coding length. Opportunity entered as the per-gene density of missense-capable coding cytosines per codon, standardized to unit standard deviation. Intervals are heteroskedasticity-robust HC1 Wald intervals, and we also assessed the interaction with a whole-gene bootstrap over 2,000 resamples, which resamples genes rather than states and therefore does not assume independence between states within a gene. We fit the model on the 4,247 genes for which every covariate is defined. The 132 genes excluded for missing coding-sequence covariates have 74 accessible of 1,274 eligible states, an accessible fraction of 5.81% versus 5.73% across the whole set. Every model estimate is reported in Supplementary Table S3b, and the per-gene model input is distributed with the reproducibility archive as gene_context_model_input.tsv.

**Expression-class construction**

We took cell-type expression classes from the Human Protein Atlas single-cell type consensus resource. For the broad classification used to define the binary post-mitotic variable, we assigned each gene a post-mitotic score and a proliferative score by summing consensus expression across the cell types in each group, then combined them as ratio = $\log_2((\text{post-mitotic} + 1) / (\text{proliferative} + 1))$. We called a gene post-mitotic-enriched at a ratio of at least 1, proliferating-enriched at a ratio of at most -1, and ubiquitous between those bounds; the thresholds correspond to a two-fold difference between the two summed scores. Genes absent from the resource received no class and were excluded from the class screens. The pseudocount of 1 gives every gene a defined ratio, so no ties required resolution and each gene falls in exactly one class.

The eight single-cell classes screened in Figure 3a are the cell types with a usable consensus-enriched gene set in the same resource, and each gene contributes to one class. We treated neurons, oligodendrocytes, photoreceptors, cardiomyocytes, and skeletal muscle as post-mitotic, and myeloid progenitors, spermatogenic cells, and basal epithelial cells as proliferative. We constructed this classification for the gene-context question after completing the accessible-state catalog, so it is not a prespecified hypothesis, and we report the analysis as exploratory.

For each class, the accessible fraction was compared with a null generated by permuting class labels across genes 2,000 times, which preserves the per-gene distribution of eligible states. The empirical two-sided P value is (b + 1) / (B + 1), where b counts permutations whose absolute deviation from the permutation mean equals or exceeds the observed absolute deviation and B = 2,000. We adjusted P values across the eight classes using the Benjamini-Hochberg procedure. The broad three-class screen used the one-sided upper-tail form of the same estimator and was adjusted across its three classes; it is reported only in Supplementary Table S3a because the binary post-mitotic variable, rather than that screen, enters the interaction model.

**Oxidized-guanine-supported sensitivity analysis**

Each of the 3,910 accessible disease-associated states carries the template-strand guanine coordinate resolved for the primary lesion-map comparison, and we called a state oxidized-guanine-supported when that coordinate was detected in at least one of the two mapping resources under the frozen mapping rules described above. We did not change the threshold for the sensitivity analysis. This gave 2,146 supported and 1,764 unsupported states, and of the 2,146 supported states, 2,118 occurred in genes included in the primary gene-context model, and 559 belonged to the higher-confidence ClinVar subset.

The joint gene-level model was refitted with the per-gene count of supported states as the binomial response, leaving the eligible-state denominator, the covariates, and the interaction term unchanged. We repeated the expression-class permutation screen on the same broad Human Protein Atlas classes, using supported states as the outcome and the same gene-label permutation

(2,000 permutations) with Benjamini-Hochberg correction across classes. The single-cell expression classes, top-level disease categories, and pathway gene sets were not retested on the supported subset, so the supported-subset result rests on the joint model and the broad expression classes.

A further model asked whether map detection itself depends on gene context. Within the 3,836 accessible states in model genes, detection was regressed on post-mitotic expression, neurological annotation, and their interaction by binomial logistic regression, adjusting for coding length, standardized C>A coding opportunity, log ClinVar record count, and the log number of accessible states per gene, with standard errors clustered by gene.

### Supplementary Figures

Supplementary Figures S1 and S2 are presented below with their legends.

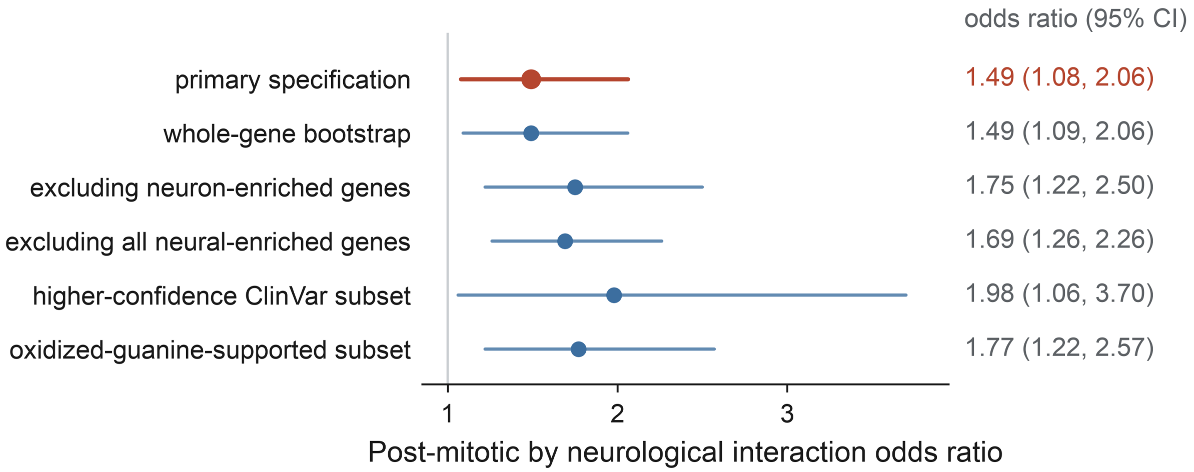


**Supplementary Figure S1.** Specification sensitivity of the gene-context interaction. Points are odds ratios for the interaction between post-mitotic expression and neurological disease annotation and horizontal bars are 95% intervals, with the primary specification in rust. The whole-gene bootstrap row uses percentile intervals and all others use heteroskedasticity-robust Wald intervals. The final row restricts accessible states to the 2,146 coordinates detected in published oxidized-guanine maps, which is a sensitivity analysis rather than a redefinition of the primary catalog, and the vertical line marks an odds ratio of 1.

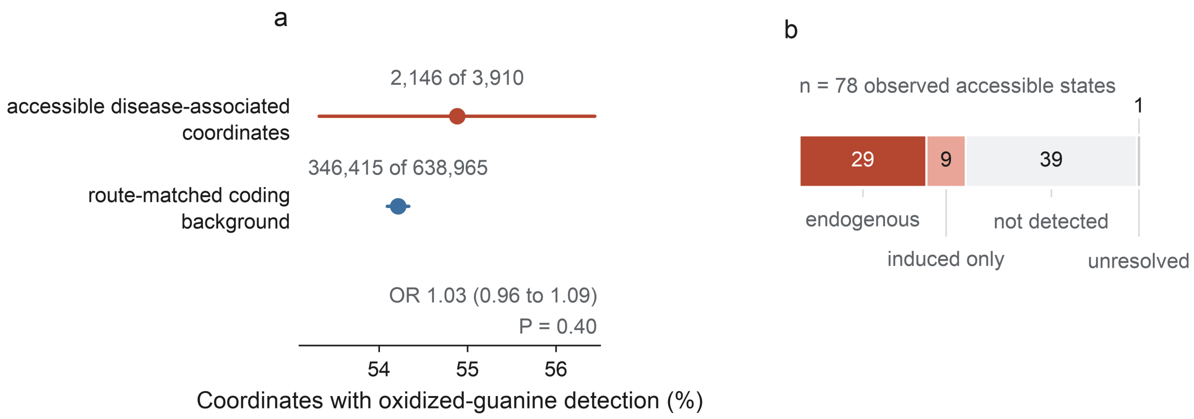


**Supplementary Figure S2.** Oxidized-guanine map evidence at accessible coordinates. (a) Proportion of coordinates detected in at least one published oxidized-guanine map, comparing accessible disease-associated coordinates with route-matched missense-capable coding cytosines in the same transcripts, with counts printed above each point; the odds ratio and its 95% confidence interval from a two-sided Fisher exact test are shown on the panel. (b) The 78 observed accessible states partitioned into the four mutually exclusive lesion-support categories, with the single unresolved state labeled above its segment.

### Supplementary Tables

The supplementary tables are provided as separate machine-readable files and indexed in the supplementary table index, which gives the table number, file name, description, row and column counts, and the result each supports.
